# Understanding the impact of preprocessing pipelines on neuroimaging cortical surface analyses

**DOI:** 10.1101/2020.05.22.100180

**Authors:** Nikhil Bhagwat, Amadou Barry, Erin W. Dickie, Shawn T. Brown, Gabriel A. Devenyi, Koji Hatano, Elizabeth DuPre, Alain Dagher, M. Mallar Chakravarty, Celia M. T. Greenwood, Bratislav Misic, David N. Kennedy, Jean-Baptiste Poline

## Abstract

The choice of preprocessing pipeline introduces variability in neuroimaging analyses that affects the reproducibility of scientific findings. Features derived from structural and functional MR imaging data are sensitive to the algorithmic or parametric differences of preprocessing tasks, such as image normalization, registration, and segmentation to name a few. Therefore it is critical to understand and potentially mitigate the cumulative biases of pipelines in order to distinguish biological effects from methodological variance. Here we use an open structural MR imaging dataset (ABIDE), supplemented with the Human Connectome Project (HCP), to highlight the impact of pipeline selection on cortical thickness measures. Specifically, we investigate the effect of 1) software tool (e.g. ANTs, CIVET, FreeSurfer), 2) cortical parcellation (DKT, Destrieux, Glasser), and 3) quality control procedure (manual, automatic). We divide our statistical analyses by 1) method type, i.e. task-free (unsupervised) versus task-driven (supervised), and 2) inference objective, i.e. neurobiological group differences versus individual prediction. Results show that software, parcellation, and quality control significantly impact task-driven neurobiological inference. Additionally, software selection strongly impacts neurobiological and individual task-free analyses, and quality control alters the performance for the individual-centric prediction tasks. This comparative performance evaluation partially explains the source of inconsistencies in neuroimaging findings. Furthermore, it underscores the need for more rigorous scientific workflows and accessible informatics resources to replicate and compare preprocessing pipelines to address the compounding problem of reproducibility in the age of large-scale, data-driven computational neuroscience.

## Introduction

Reproducibility, a presumed requisite of any scientific experiment, has recently been under scrutiny in the field of computational neuroscience [1–7]. Specifically, replicability and generalizability of several neuroimaging pipelines and the subsequent statistical analyses have been questioned, potentially due to insufficient sample size [8], imprecise or flexible methodological and statistical apriori assumptions [9–11], and poor data/code sharing practices [12,13]. Broadly speaking, reproducibility can be divided in two computational goals [14]. The first goal is replicability, which implies that a re-executed analysis on the identical data should always yield the same results. The second goal pertains to generalizability, which is assessed by comparing the scientific findings under variations of data and analytic methods. Typically, findings are deemed generalizable when similar (yet independent) data and analysis consistently support the experimental hypothesis. This in turn raises the issue of defining what constitutes “similar” data and analytic methodology. Nonetheless, traditionally experimental validation on independent datasets has been utilized to assess generalizability. However, as the use of complex computational pipelines has become an integral part of modern neuroimaging analysis [15], comparative assessment of these pipelines and their impact on the generalizability of findings deserves more attention.

We present a comparative assessment of multiple structural neuroimaging preprocessing pipelines on the Autism Brain Imaging Data Exchange (ABIDE), a publicly accessible dataset comprising healthy controls and individuals with autism spectrum disorder (ASD) [18]. A few studies have previously highlighted the variability in neuroimaging analyses introduced by the choice of a preprocessing pipeline for structural MR images [16,17], however they have not focused on the relative impact of analysis tools, quality control, and parcellations on the consistency of results. The inconsistencies in the results arise from several algorithmic and parametric differences that exist in the preprocessing tasks, such as image normalization, registration, segmentation, etc. within pipelines. It is critical to understand and potentially mitigate the cumulative biases of the pipelines to disambiguate biological effect from methodological variance. We further replicate our findings on the Human Connectome Project (HCP) data.

For this purpose, we propose a comprehensive investigation of the impact of pipeline selection on cortical thickness measures, a widely used (3129 hits on PubMed and 42,200 hits on Google Scholar for “cortical thickness” AND “Magnetic resonance imaging” search query), fundamental phenotype, and its statistical association with biological age. We limit the scope of pipeline variation to three axes of parameter selection: 1) image processing tool, 2) anatomical priors, 3) quality control (see Fig 1). The impact of the variation is measured on two types of statistical analyses, namely: 1) neurobiological inference carried out using general linear modeling (GLM) techniques at the group level; and 2) individual predictions from machine-learning (ML) models. We note that here the focus is on the preprocessing stages of a computational pipeline, and the impact of dataset and statistical model selection is thus out of the current scope. Our goal is not to explain potential differences in results or establish criteria to rank pipelines or tools, but to document the pipeline effect and provide best practice recommendations to the neuroscience community with respect to pipeline variation, also referred to as pipeline vibration effects.

**Fig. 1.**
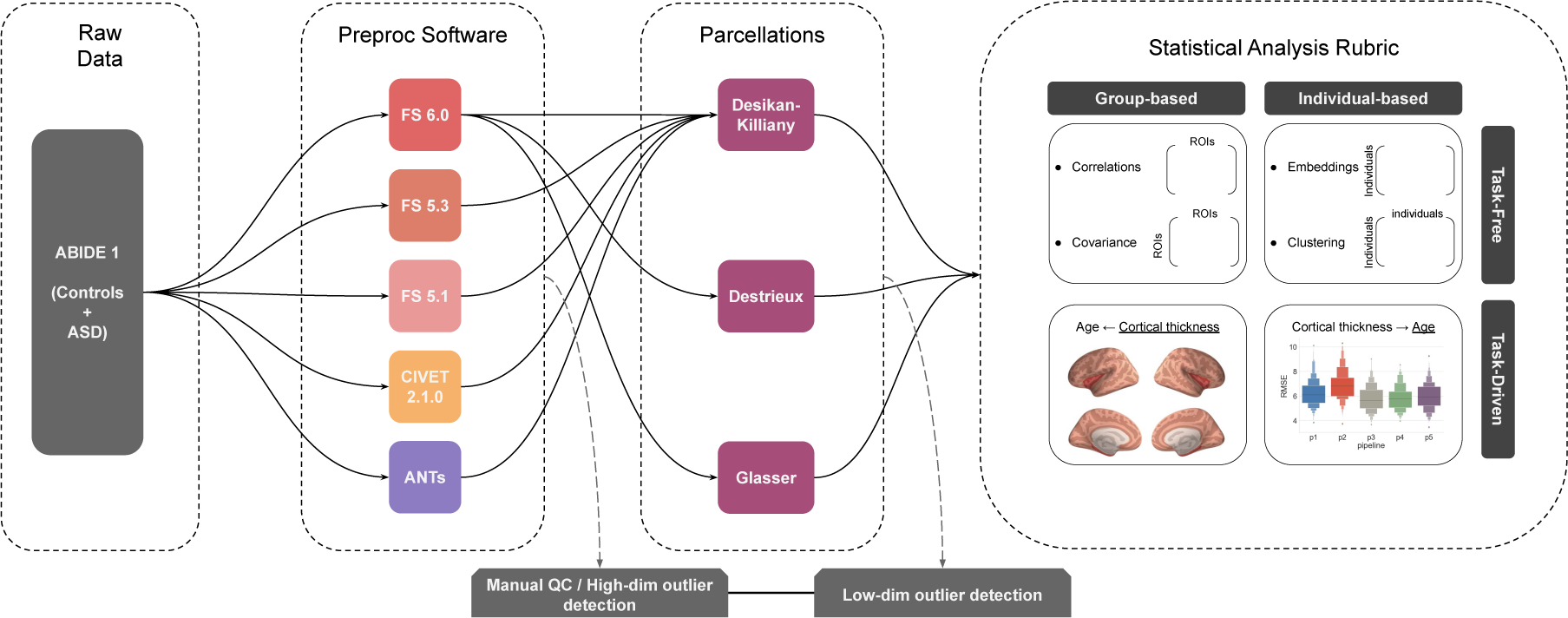
Preprocessing pipeline building blocks and potential permutations for a typical structural MR image analysis. Only a subset of the possible pipelines is analyzed and shown with arrows. Note that manual quality control and automatic outlier detection can be performed at various stages.

Although here we do not focus on identifying biological differences between ASD case and control groups, we use the case-control samples to gain insight into the effect of diagnosis on reproducibility analysis - which is a critical evaluation for clinical applications. Additionally, we use a data sample from the HCP as a validation dataset (Van Essen DC et al. 2013) to assess if our findings replicate on an independent dataset. Note that the scope of this secondary analysis is limited to a proof of concept dataset comparison.

We organize our comparative assessments on the ABIDE dataset as follows. We report comparisons across the three aforementioned axes of variation. This comprises five neuroimaging preprocessing tools: 1) FreeSurfer 5.1, 2) FreeSurfer 5.3, 3) FreeSurfer 6.0, 4) CIVET 2.1.0, and 5) ANTs; three anatomical priors (i.e. cortical parcellations): 1) Desikan-Killiany-Tourville, 2) Destrieux, and 3) Glasser; and five quality control (QC) procedures 1) No QC 2) manual lenient 3) manual stringent, 4) low-dimensional automatic outlier detection (i.e. <500 ROIs), and 5) high-dimensional automatic outlier detection (i.e. > 100k vertices). The entire combinatorial set of comparisons (5 software x 3 parcellations x 5 QC) is not feasible due to practical limitations (described later), and therefore we report results for five tools procedures and three atlases across five quality control procedures (5 software + 3 parcellations) x 5 QC, as shown by the connecting arrows in Fig 1. We use these 40 preprocessed data with four types of statistical analyses based on a method type (i.e. task-free vs. task-driven) and an inference objective (neurobiological vs. individual), as described in detail in the methods.

## Materials and Methods

### Participants

Participants from the ABIDE dataset were used for this study [18]. The ABIDE 1 dataset comprises 573 control and 539 autism spectrum disorder (ASD) individuals from 16 international sites. The neuroimaging data of these individuals were obtained from the ABIDE preprocessing project [19], the Neuroimaging Tools and Resources Collaboratory (NITRC) (http://fcon_1000.projects.nitrc.org/indi/abide/abide_I.html), and the DataLad repository (http://datasets.datalad.org/?dir=/abide/RawDataBIDS). Different subsets of individuals were used for various analyses based on 1) specific image processing failures, 2) need for a common sample set for software tool comparison, and 3) quality control procedures. The demographic description of these subsets is provided in Table 1, and Figure 2. The complete lists of subjects can be obtained from the code repo: https://github.com/neurodatascience/compare-surf-tools

**Table 1.** Subject demographics for different analyses.

| Comparisons | QC | Diagnosis | Subjects (N) | Age (mean, sd) | Sex (M/F) |
| --- | --- | --- | --- | --- | --- |
| Software tools | No QC(N=778) | Controls | 415 | 17.8, 7.7 | 346/69 |
|  |  | ASD | 363 | 18.3, 8.7 | 320/43 |
|  | Lenient Manual(N=748) | Controls | 407 | 17.8, 7.6 | 338/69 |
|  |  | ASD | 341 | 18.4, 8.8 | 300/41 |
|  | Stringent Manual(N=194) | Control | 113 | 15.6, 5.5 | 93/20 |
|  |  | ASD | 81 | 16.2, 5.8 | 71/10 |
|  | Auto QC low-dim(N=683) | Controls | 371 | 16.2, 5.4 | 309/62 |
|  |  | ASD | 312 | 15.9, 5.0 | 276/36 |
|  | Auto QC high-dim(N=662) | Controls | 356 | 15.6, 5.0 | 293/63 |
|  |  | ASD | 306 | 15.7, 4.9 | 269/37 |
| Parcellations | No QC(N=1047) | Controls | 552 | 17.0, 7.5 | 456/96 |
|  |  | ASD | 495 | 17.1, 8.4 | 436/59 |
|  | Lenient Manual(N=975) | Controls | 525 | 17.1, 7.5 | 430/95 |
|  |  | ASD | 450 | 17.4, 8.6 | 395/55 |
|  | Stringent Manual(N=240) | Controls | 137 | 15.0, 5.6 | 112/25 |
|  |  | ASD | 103 | 16.1, 6.3 | 91/12 |
|  | Auto QC low-dim(N=961) | Controls | 516 | 15.6, 5.6 | 422/94 |
|  |  | ASD | 445 | 15.0, 5.1 | 390/55 |
|  | Auto QC high-dim(N=912) | Controls | 483 | 15.0, 4.9 | 393/90 |
|  |  | ASD | 429 | 14.9, 4.9 | 377/52 |

**Fig. 2.**
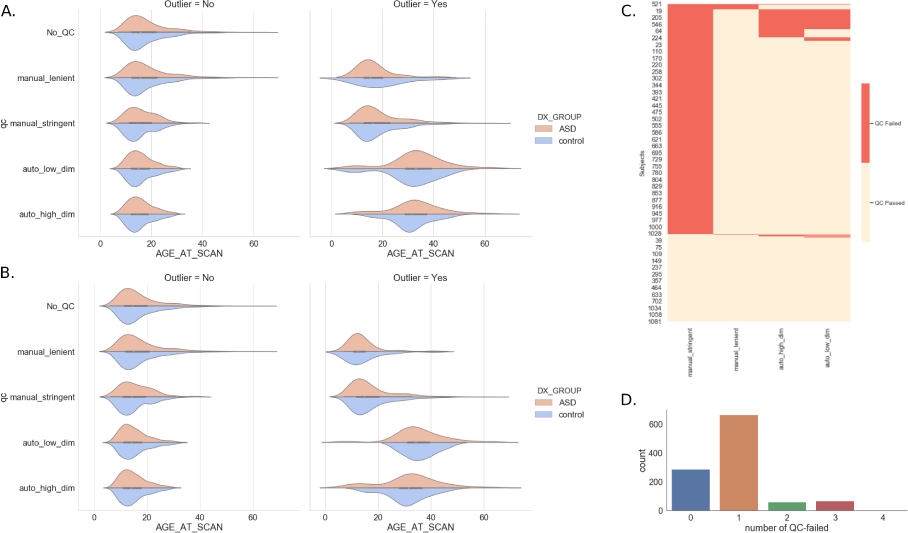
Age distributions for sample subsets used for (A) software comparison and (B) parcellation comparison analyses. See Table 1 for sample sizes. Failed QC overlap across manual QC and automatic outlier detection procedures is show in (C). Distribution of total outlier count (sum) based on four possible manual QC and automatic outlier detection procedures is shown in (D)

### MR Image processing and cortical thickness measurements

#### FreeSurfer

FreeSurfer (FS) delineates the cortical surface from a given MR scan and quantifies thickness measurements on this surface for each brain hemisphere [20,21]. The default pipeline consists of 1) affine registration to the MNI305 space [22]; 2) bias field correction; 3) removal of skull, cerebellum, and brainstem regions from the MR image; 3) estimation of white matter surface based on MR image intensity gradients between the white and grey matter; and 4) estimation of pial surface based on intensity gradients between the grey matter and cerebrospinal fluid (CSF). The distance between the white and pial surfaces provides the thickness estimate at a given location of cortex. For detailed description refer to [23]. The individual cortical surfaces are then projected onto a common space (i.e. fsaverage) characterized by 163,842 vertices per hemisphere to establish inter-individual correspondence.

In this work, the cortical thickness for each MR image was computed using FS 5.1, 5.3, and 6.0 versions. The FS5.1 measurements were obtained from the ABIDE preprocessing project [19]. Standard recon-all pipeline with “-qcache” flag was used to process and resample the images onto common (fsaverage) space. The FS5.3 measurements were extracted using the standard ENIGMA cortical thickness pipeline [24]. Lastly, the FS6.0 measurements were obtained using the standard recon-all pipeline with “-qcache” flag as well. Compute Canada [25] and CBRAIN [26] computing infrastructures were used for processing of FS5.3 and FS6.0 data.

#### CIVET

CIVET 2.1 (http://www.bic.mni.mcgill.ca/ServicesSoftware/CIVET-2-1-0-Introduction) preprocessing was performed on the data obtained from NITRC. The standard CIVET pipeline consists of 1) N3 bias correction [27]; 2) affine registration to the MNI ICBM 152 stereotaxic space; 3) tissue classification into white matter (WM), grey matter (GM) and cerebrospinal fluid; 4) brain splitting into left and right hemispheres for independent surface extraction; 5) estimation of WM, pial, and GM surfaces. The cortical thickness is then computed using the distance (i.e. Tlink metric) between WM and GM surfaces at 40,962 vertices per hemisphere.

#### ANTs

The MR imaging dataset preprocessed with ANTs (“RRID:SCR_004757, version May-2017”) was obtained from the ABIDE preprocessing project [19]. The detailed description of ANTs cortical thickness pipeline can be found here [16]. Briefly, the ANTs pipeline consists of 1) N4 bias correction [28]; 2) brain extraction; 3) prior-based segmentation and tissue-based bias correction; and 4) Diffeomorphic registration-based cortical thickness estimation [29].

One key differentiating aspect of ANTs is that it employs quantification of cortical thickness in the voxel-space, unlike FreeSurfer or CIVET, which operate with vertex-meshes.

### Cortical parcellations

The regions of interest (ROI) were derived using three commonly used cortical parcellations, namely 1) Desikan-Killiany-Tourville (DKT) [30], 2) Destrieux [31], and 3) Glasser [32]. DKT parcellation consists of 31 ROIs per hemisphere and is a modification of the Desikan–Killiany protocol [33]) to improve cortical labeling consistency. DKT label definitions are included in all three FreeSurfer (FS), CIVET, and ANTs pipelines, which allows the comparison of cortical phenotypic measures across these tools. The Destrieux parcellation is a more detailed anatomical parcellation proposed for a precise definition of cortical gyri and sulci. The Destrieux parcellation comprises 74 ROIs per hemisphere, and is also available in the FS pipeline. In contrast to these structural approaches, the Glasser parcellation was created using multimodal MR acquisitions from 210 HCP subjects [34] with 180 ROIs per hemisphere. Glasser label definitions are available in the “fsaverage” space (https://doi.org/10.6084/m9.figshare.3498446.v2), i.e. the common reference space used by FreeSurfer, allowing comparisons across multiple parcellations.

### Quality Control

We employed manual (i.e. visual) and automatic (statistical outlier detection) procedures to investigate the effect of quality control (QC) on thickness distributions derived from combinations of the different software tools and cortical parcellations. The manual quality checks were performed on the extracted cortical surfaces by two independent expert raters [35,36]. The two raters used different criteria for assessing the quality of surface delineation. This in turn yielded two lists of QC-passed subjects from “lenient” and “stringent” criteria. We note that these lenient and stringent QC lists were generated independently using FS and CIVET images, respectively; and then applied to all pipeline variations. The automatic quality control was performed using an outlier detection algorithm based on a random min-max multiple deletion (RMMMD) procedure (Barry et al. in preparation). The RMMMD algorithm is a high-dimensional extension of Cook’s influence measure to identify influential observations. The outlier detection method was applied separately to high-dimensional vertex-wise output and low-dimensional aggregate output based on cortical parcellations for each software and parcellation choice.

### Statistical Analysis

We categorize the downstream statistical analyses into a 2×2 design. The first factor consists of either 1) unsupervised, task-free (TF) analyses or 2) supervised, task-driven (TD) analyses. The second factor corresponds to either 1) neurobiological (N) tasks investigating the biological effect across groups of individuals or 2) individual (I) tasks predicting individual-specific states (see Table 2). The task-free, neurobiologically oriented analyses (TF-N) aim at quantifying similarity of preprocessed features (i.e. ROI-wise cortical thickness values) without the explicit constraint of an objective function. Task-driven, neurobiologically oriented analyses (TD-N) quantify feature similarity in the context of a general linear model (GLM) framework. Individually oriented analyses formulate the duality of neurobiological analyses, with a focus on individual similarity in task-free (TF-I) and task-driven (TD-I) contexts.

**Table 2.**
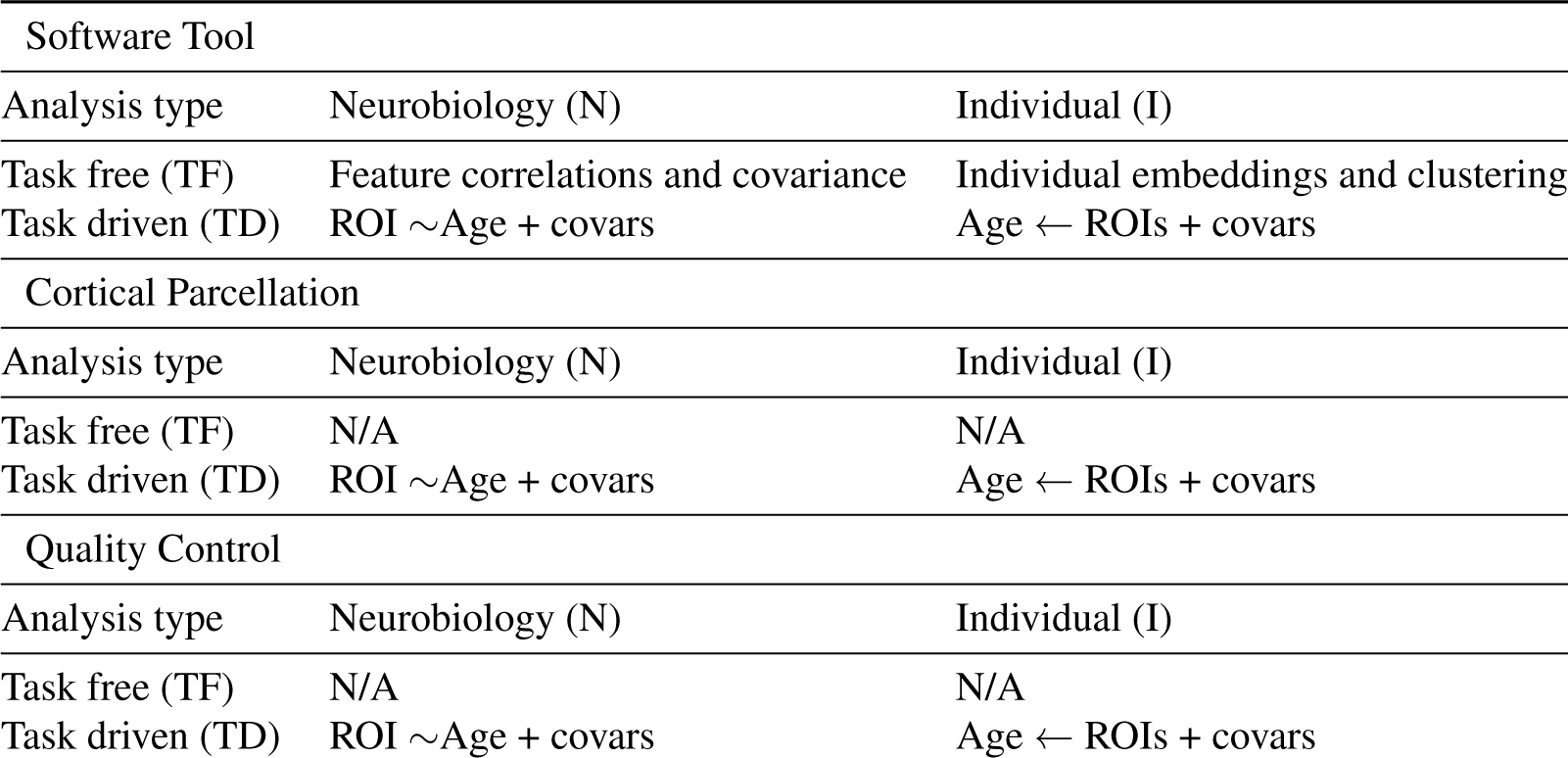
2×2 rubric showing types of analysis performed for each axis of variation.

Previous work has reported varying degrees of association and predictability of age from cortical thickness measures in neurotypical and ASD cohorts [37–41]. We therefore selected biological age as our objective for the task-driven (TD) analyses. Although other clinical variables (e.g. diagnosis) could be used, availability and unambiguity of age quantification across datasets simplifies comparison of the different analyses.

For TF-N analysis we evaluate the pairwise correlation and covariance of features using Pearson’s r metric. For TF-I analysis, we assess individual similarity using t-SNE and hierarchical clustering with Euclidean distance and Ward’s linkage metrics. For TD-N analysis, we build a GLM to associate cortical thickness and biological age with sex and data collection site as covariates. For TD-I analysis, we train a random forest (RF) model for age prediction using cortical thickness, sex, and data collection site as predictors. Of note, we also assess the importance assigned to cortical features by the RF model. Machine learning (ML) model performance and feature importance is assessed within 100 iterations of a shuffle-split cross-validation paradigm.

We also note that not all pipeline variations can be assessed easily within this to 2×2 statistical analyses design. As mentioned before we only analyze a subset ((5+3)x5) of possible pipeline variations, and compare the five software tools using common DKT parcellation. Tool comparison with Destrieux and Glasser parcellations is not trivial due to their unavailability for CIVET and ANTs. This also limits our comparison across three parcellations solely with FreeSurfer 6.0. We do however compare all five QC procedures with these combinations. The analyses performed in this work are provided in Table 2. The code used for the analyses is available here: https://github.com/neurodatascience/compare-surf-tools.

### Validation Study

The T1w images of 1108 individuals from the HCP dataset [42] were successfully preprocessed using FS 6.0 and CIVET 2.1 respectively, and average cortical thickness measurements in the DKT ROIs were obtained. Identical to the ABIDE analysis, we evaluated the pairwise correlations and covariance of features between CIVET 2.1 and FS 6.0 using Pearson’s r metric, then we compared it using the same approach as for the ABIDE dataset.

## Results

### Task-free neurobiological (TF-N) analysis

Feature comparisons across the five software tools are performed using common DKT parcellation. The pairwise comparisons between software tools are performed based on the ROI-wise Pearson correlations between thickness measures produced by each tool (See Figure 3, Table 3). The pairwise comparisons between FS, CIVET, and ANTs tools show very little similarity with correlation values averaged over all regions remaining low (*rϵ*[0.39, 0.52]). The comparisons between different versions of FS show relatively better average correlation performance (*rϵ*[0.83, 0.89]). Stratifying comparisons by diagnosis does not improve correlation. ROI specific performance shows the lowest median correlation for the left rostral-anterior-cingulate (r=0.27), left and right isthmus-cingulate (r=0.29,0.31) regions, and the highest median correlation for the left cuneus (r=0.63), right postcentral (r=0.63), and left caudal-middle-frontal (r=0.62) regions across all software pairs. The pairwise thickness distributions for three randomly selected exemplar ROIs corresponding to different levels of median correlations across software tools are shown in Figure 3. The exemplar ROI comparison suggests that ROIs with high correlation levels tend to have lower overlap between the pairwise thickness distributions.

**Table 3.** Average ROI correlations between software pairs for control and ASD cohorts.

|  | Controls |  |  |  |  | ASD |  |  |  |  |
| --- | --- | --- | --- | --- | --- | --- | --- | --- | --- | --- |
|  | ANTs | CIVET | FS5.1 | FS5.3 | FS6.0 | ANTs | CIVET | FS5.1 | FS5.3 | FS6.0 |
| ANTs | 1 | 0.43 | 0.45 | 0.48 | 0.44 | 1 | 0.39 | 0.39 | 0.46 | 0.41 |
| CIVET |  | 1 | 0.48 | 0.52 | 0.52 |  | 1 | 0.44 | 0.48 | 0.49 |
| FS5.1 |  |  | 1 | 0.89 | 0.84 |  |  | 1 | 0.87 | 0.83 |
| FS5.3 |  |  |  | 1 | 0.89 |  |  |  | 1 | 0.88 |
| FS6.0 |  |  |  |  | 1 |  |  |  |  | 1 |

**Fig. 3.**
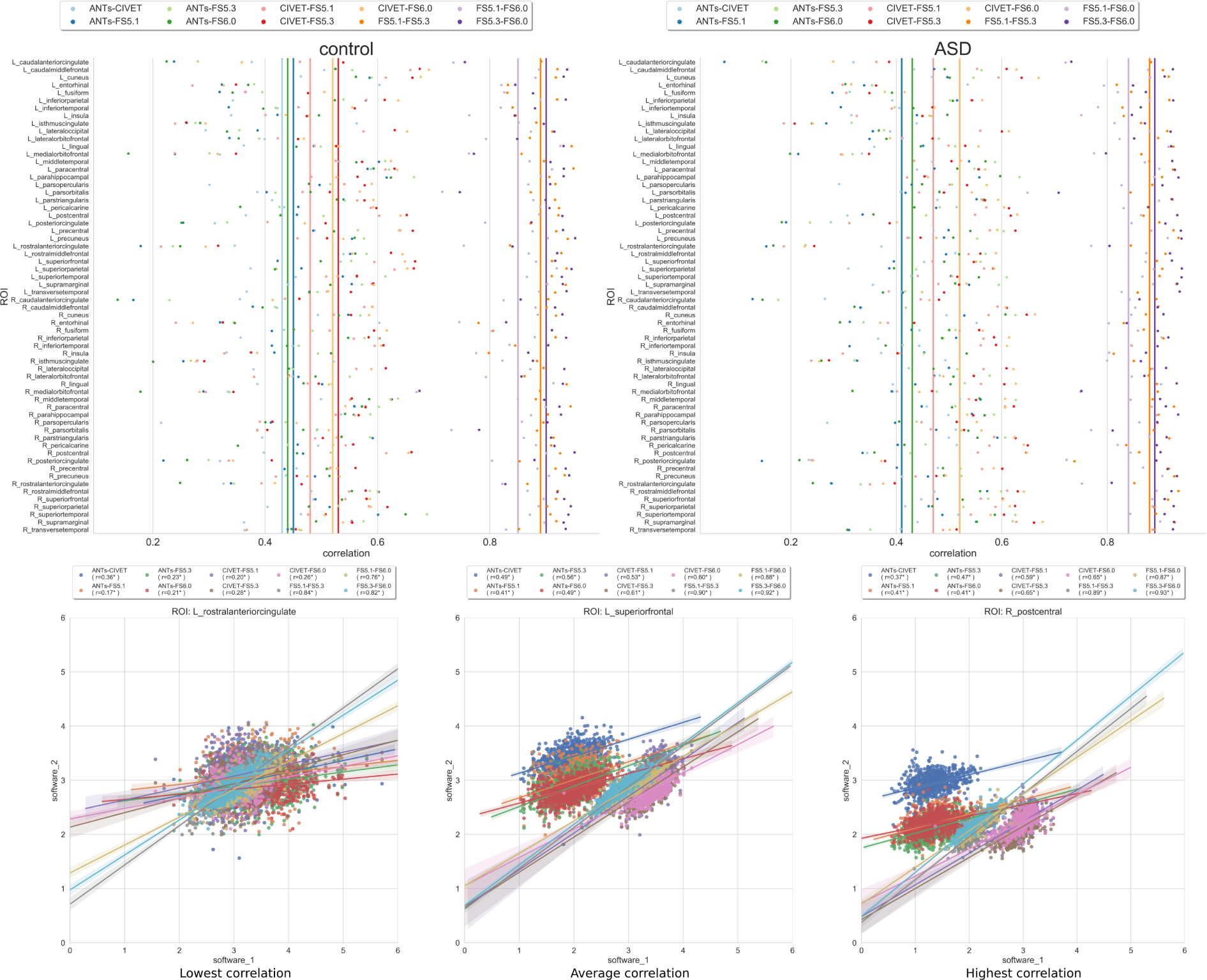
Task Free - Neurobiology (TF-N) analysis: Top) Correlation between cortical thickness values for software pairs measured independently over ROIs for control and ASD groups. The vertical lines represent the mean correlation across all ROIs. The ROIs are defined using Desikan-Killiany-Tourville (DKT) parcellation. Bottom) Distribution of cortical thickness values for exemplar ROIs with lowest, average, and highest median correlation across software pairs.

The covariance matrix of ROIs and subsequently derived structural network metrics reveal several software specific differences. First, the covariance matrix shows large variation of patterns across software tools (see Figure 4-middle). All software tools show strong bilateral symmetry evidenced by the high correlation values on the diagonal representing hemispheric ROI pairs. Interestingly, CIVET features show stronger intra-hemispheric correlation between ROIs compared to the inter-hemispheric values. The DKT ROIs are grouped based on their membership in the Yeo resting state networks [43] to compute graph theoretic metrics. Figure 4 shows the variation in the two commonly used metrics. Figure 4-top shows the impact of correlation threshold, typically used for denoising graph-edges, on the fundamental measure of graph density. The three FS versions show relatively similar performance for all resting state networks, with somato-motor and default mode exhibiting highest and lowest densities, respectively. Compared to FS values, ANTs and CIVET show different magnitudes and/or rankings of graph densities across networks. These differences are further amplified in the graph degree-centrality measurements across networks.

**Fig. 4.**
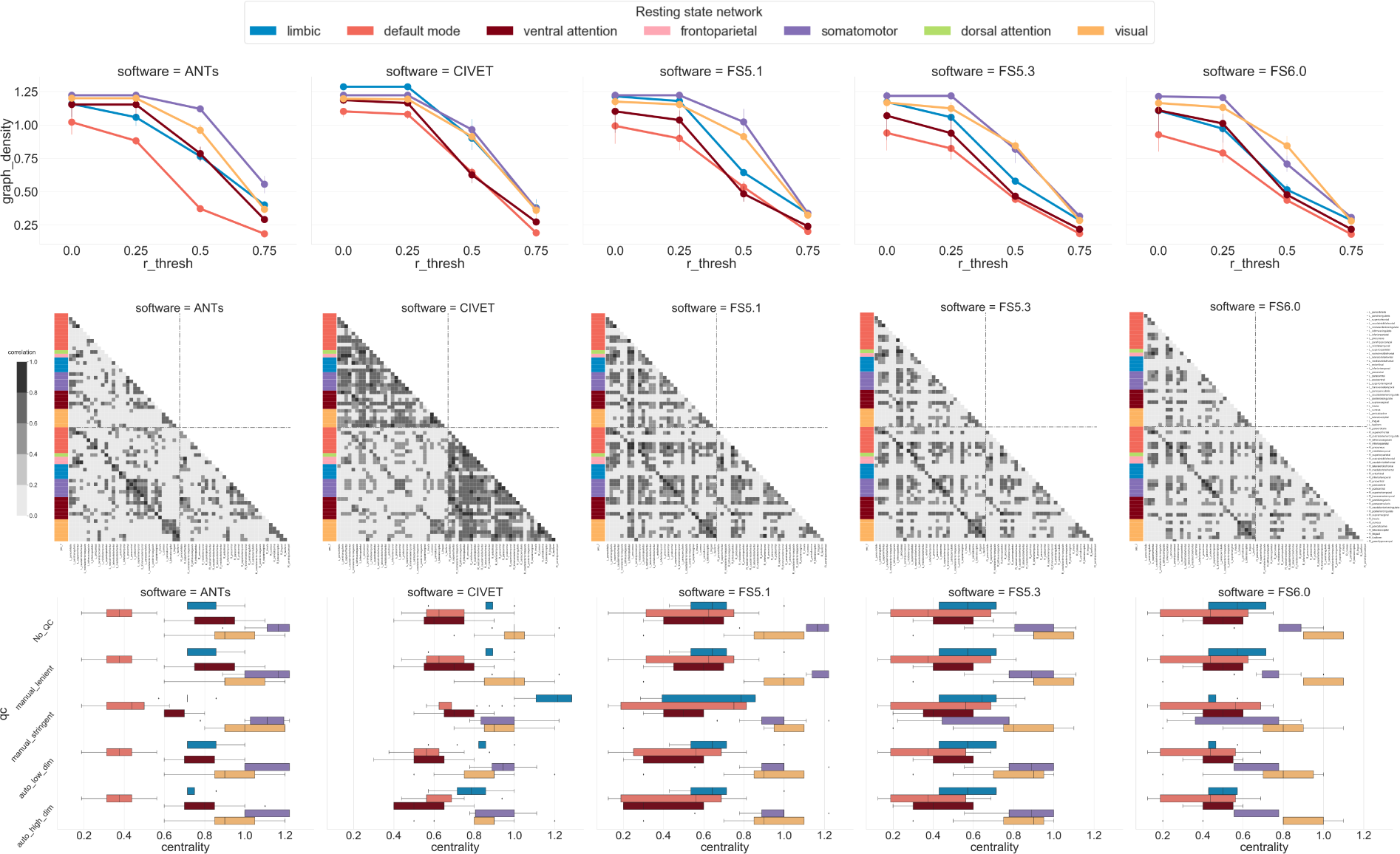
Task-Free neurobiological (TF-N) analysis: Top) Graph density for different correlation cutoff thresholds used for constructing a structural network. The error bars show variation due to the QC procedure. Middle) Structural covariance of each software measured as inter-ROI correlation with cutoff value of 0.5. For simplicity, the covariance plot is generated with original data. The covariance patterns are grouped by Yeo resting state networks membership. Bottom) Distribution of regional degree-centrality metric per Yeo network for each software with different QC procedures. Note that fronto-parietal and dorsal attentional networks are excluded from some analyses due to the small number of DKT ROIs in these networks.

Figure 4-bottom shows high intra-network regional variance in degree-centrality for FS versions. This variance is relatively smaller for ANTs and CIVET but these software show largely different magnitudes of centrality, particularly in limbic and default mode networks.

Comparison across QC procedures did not show any substantial impact on correlation values. Feature comparison for a given software tool (e.g. FS6.0) across different parcellations is not trivial due to the lack of correspondence between various parcellation schemes.

### Task-free individual (TF-I) analysis

Individual comparisons using thickness measures from DKT parcellation are performed across the five software tools with an identical set of subjects. Commonly used 2-dimensional t-SNE embeddings show strong similarity between subjects for a given software tool (see Figure 5). The three FS versions are much more similar to each other than any FS version is to CIVET or ANTs, reflecting that the different versions of FS share methodological and technical components. Individual covariance is quantified using clustering consistency (CC) that measures the fraction of pairs of individuals assigned to the same cluster with two different feature sets (e.g. ANTs vs. CIVET). Based on CC metric, hierarchical clustering with Euclidean distance similarity and Ward’s linkage criterion shows poor stability (*CCϵ*[0.52, 0.61]) across software tools and between FS versions (see Table 4). In contrast, hierarchical clustering with correlation metric and average linkage criterion shows highly stable cluster membership (*CCϵ* [0.962, 0.997]).

**Table 4.** Clustering consistency between software pairs. The diagonal shows expected overlap based on 100 bootstrap samplings of features (31 ROIs) for a given software tool.

|  | Similarity: Euclidean distance, linkage: Ward's method |  |  |  |  | Similarity: correlation, linkage: average |  |  |  |  |
| --- | --- | --- | --- | --- | --- | --- | --- | --- | --- | --- |
|  | ANTs | CIVET | FS5.1 | FS5.3 | FS6.0 | ANTs | CIVET | FS5.1 | FS5.3 | FS6.0 |
| ANTs | 0.797 | 0.5 | 0.521 | 0.517 | 0.522 | 0.991 | 0.970 | 0.962 | 0.972 | 0.972 |
| CIVET |  | 0.717 | 0.5 | 0.5 | 0.5 |  | 0.994 | 0.982 | 0.992 | 0.992 |
| FS5.1 |  |  | 0.78 | 0.609 | 0.529 |  |  | 0.997 | 0.990 | 0.985 |
| FS5.3 |  |  |  | 0.703 | 0.499 |  |  |  | 0.997 | 0.995 |
| FS6.0 |  |  |  |  | 0.619 |  |  |  |  | 0.997 |

**Fig. 5.**
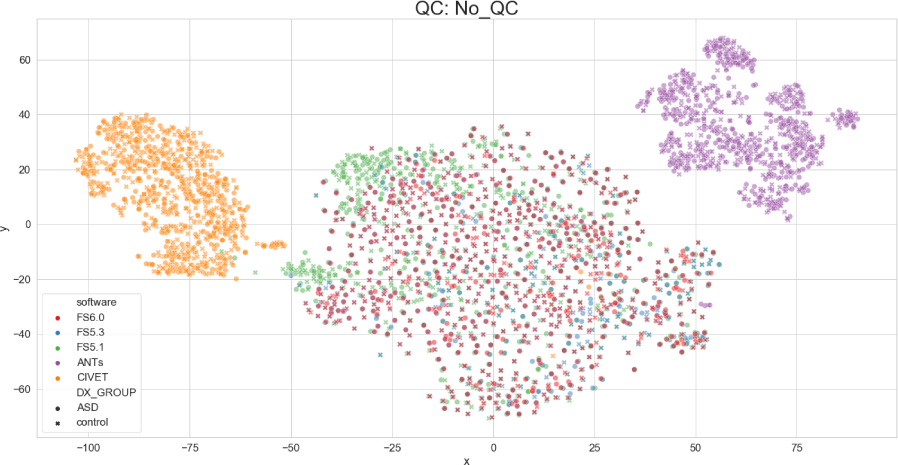
Task-free individual (TF-I) analysis: Two dimensional t-SNE representation of all individuals (No QC). The colors indicate the software tool used, and the marker style refers to the diagnostic group.

Comparison across QC procedures did not show any substantial impact on t-SNE representations or clustering consistency values. Individual comparisons across different parcellations for a given software tool (e.g. FS6.0) are not particularly informative due to the lack of correspondence between various parcellation spaces.

### Task-driven neurobiological (TD-N) analysis

The mass-univariate regression models per ROI region suggest cortex-wide association between age and thickness values for all software tools, with the exception of the CIVET-based analysis, which excludes bilateral insular regions (see Figure 6). QC procedures seem to have varying impact on the significant regions depending on the software tool. The aggregate ranking suggests higher variation in significant regions for ANTs and CIVET. In contrast the FreeSurfer versions offer relatively similar performance - with consistent exclusion of entorhinal regions. The stringent manual QC sample severely reduces the number of significant regions, which may be due to reduced statistical power.

**Fig. 6.**
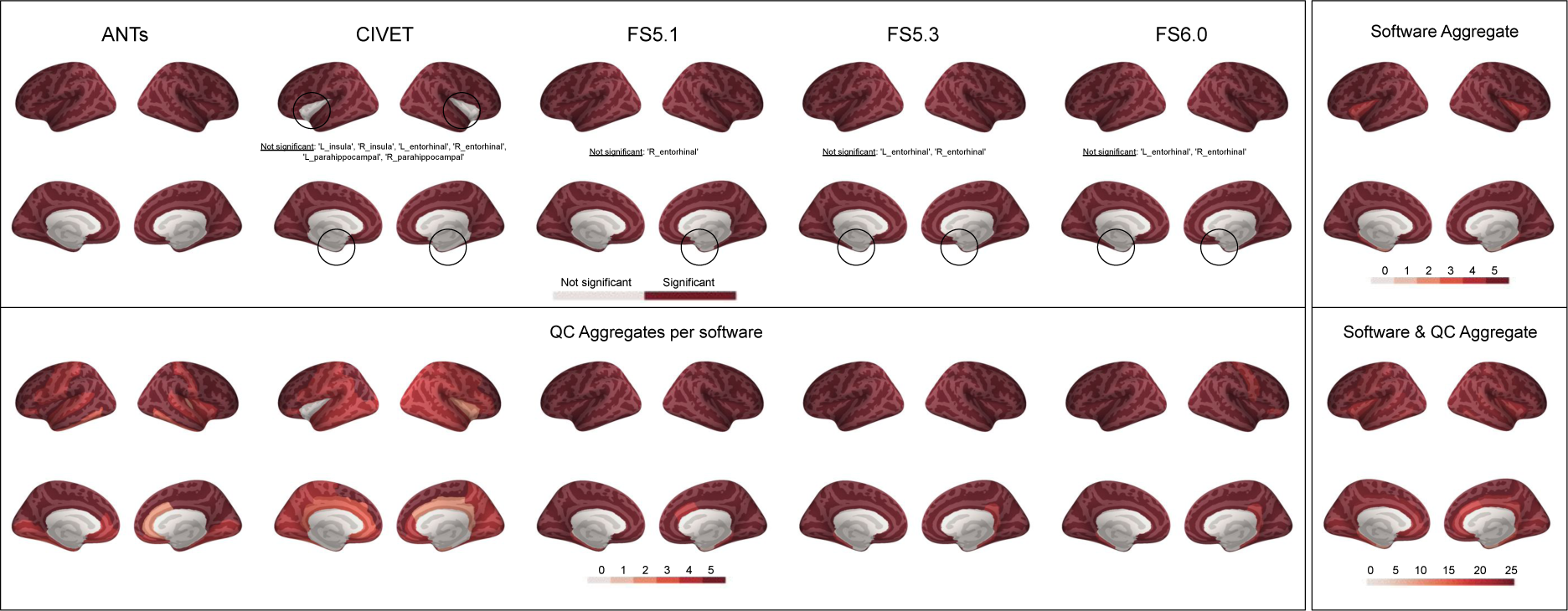
TD-N analysis: Significant ROI differences with various software and QC levels. Significance levels are corrected for multiple comparisons. Aggregate ranks are assigned based on performance agreement among five software and five QC procedures. Lower rank implies fewer QC procedures yielding the same results.

Parcellation comparisons for FreeSurfer 6.0 reaffirm cortex-wide association between age and thickness values across the three parcellation schemes with some exclusions in medial and superior temporal gyri for Destrieux and STGa, PIR, TGd, TGv, PHA1, EC, PeEc with Glasser (see Figure 7). Lenient QC does not seem to change the distribution of significant regions. However, stringent and automatic QC based results additionally exclude regions from precentral gyri for all three atlases.

**Fig. 7.**
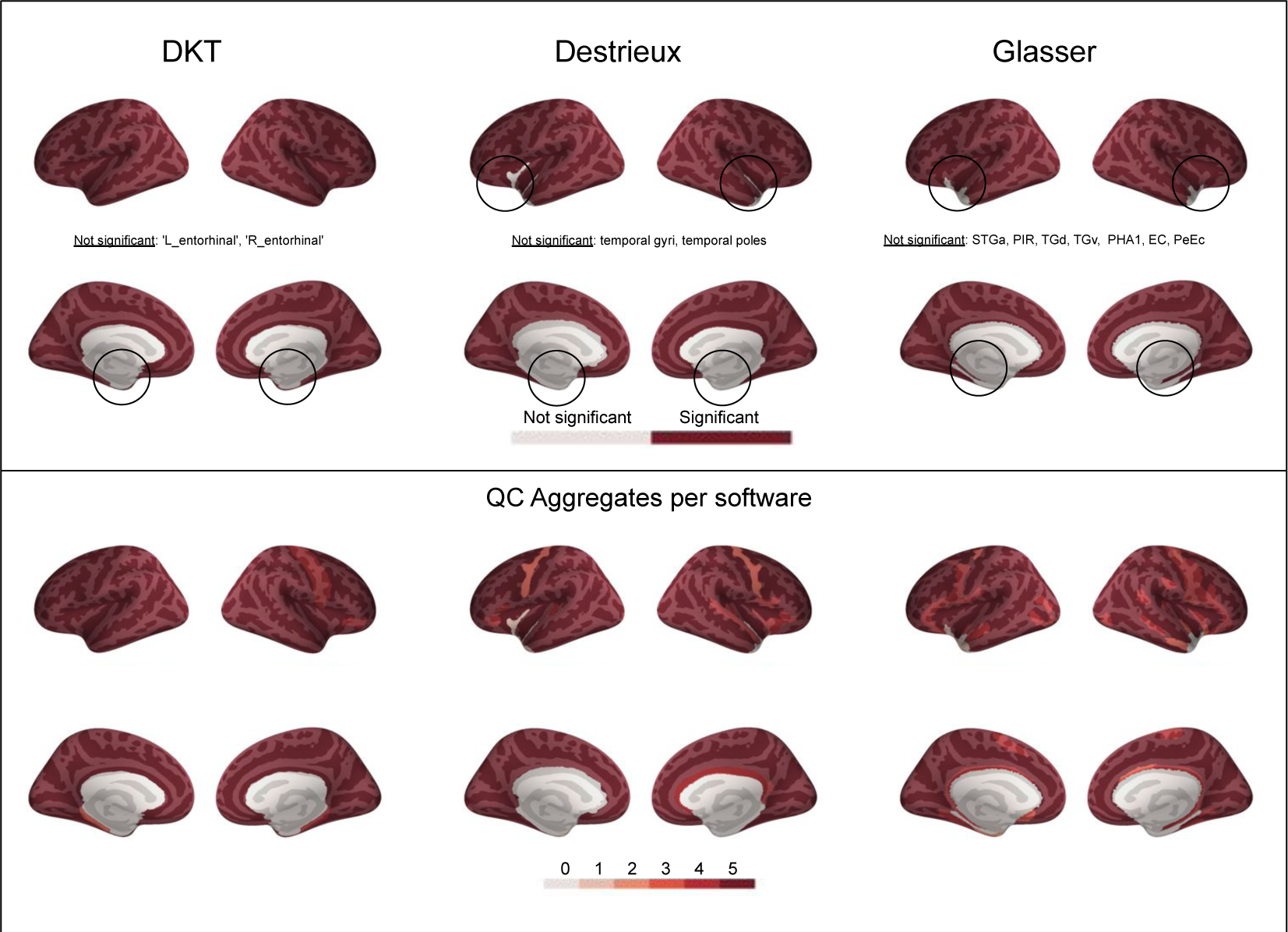
TD-N analysis: Significant ROI differences with various parcellations and QC levels. Significance levels are corrected for multiple comparisons using the Bonferoni procedure. Aggregate ranks are assigned based on performance agreement between the five QC procedures. Lower rank implies fewer QC procedures yielding the same results.

### Task-driven individual (TD-I) analysis

The RF model based predictions show consistent Root Mean Square Error (RMSE) performance (5.7 - 7.2 years) across software tools, with FS versions showing marginally lower error (see Figure 8). All model performances are statistically significant when compared against a null model. The average RMSE for the control cohort is lower than the ASD cohort; as expected per the null model, however the difference is statistically insignificant. Lenient QC does not have an impact on RMSE distributions. Stringent QC reduces the average RMSE for all software tools (3 - 5 years) and the null model. Automatic QC reduces the average RMSE as well as its variance for all software tools (3.8 - 4.7 years). Interestingly with the automatic QCs (low- and high-dimensional), the null models expectations are reversed as the average RMSE for ASD subjects is now lower than that of controls.

**Fig. 8.**
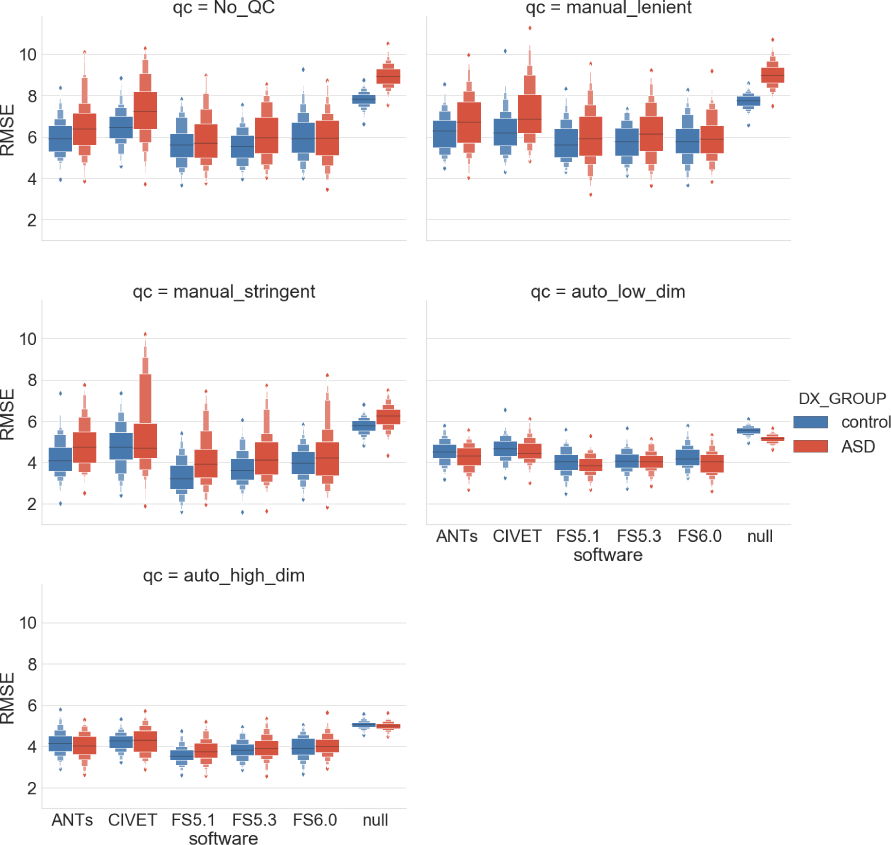
Task-driven individual (TD-I) analysis: Individual age prediction with various software and QC levels stratified by diagnosis. Performance is cross-validated using a Random Forest model over 100 shuffle-split iterations.

Parcellation-based comparisons show similar RMSE performance despite the differences in granularity of regions and the consequent number of input features to the ML models (see Figure. 9). The RMSE trends with respect to QC are also consistent, with both stringent and automatic QC reducing the average RMSE and the latter yielding a much tighter distribution of error. The null model shows lower expected error for the control cohort compared to the ASD, except for the automatic QC based analyses, where this expectation is reversed.

**Fig. 9.**
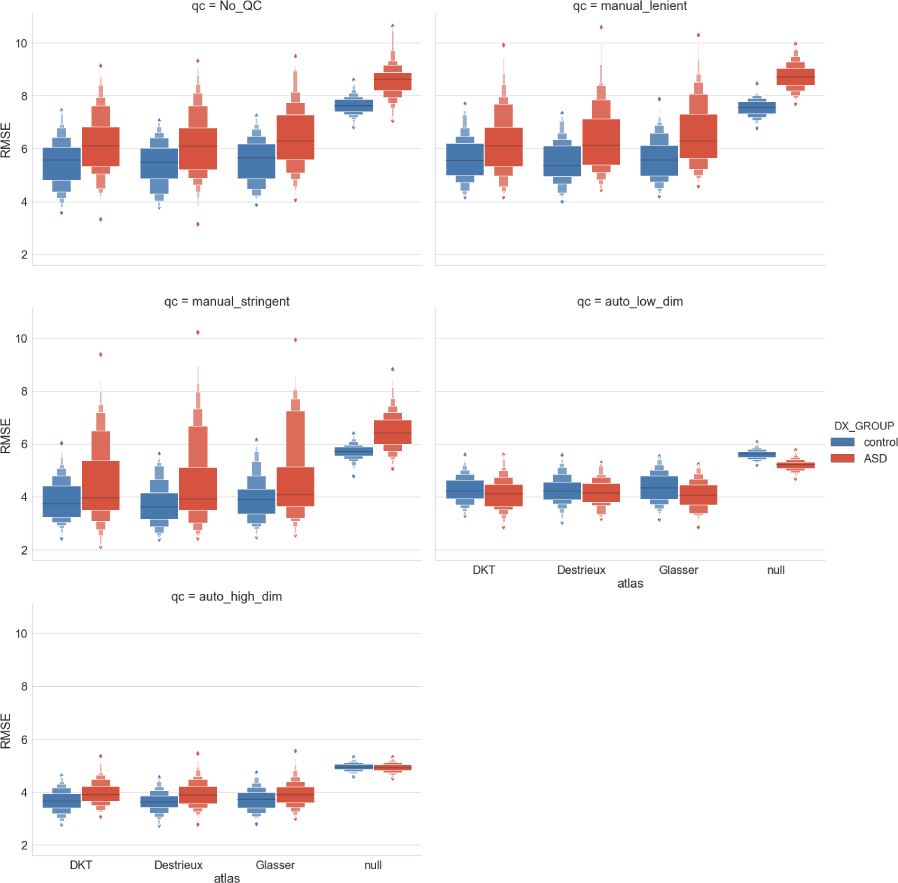
Task-driven individual (TD-I) analysis: Individual age prediction with various parcellations and QC levels stratified by diagnosis. Performance is cross-validated using a Random Forest model over 100 shuffle-split iterations.

### ROI importance from Random Forest (RF)

The cross-validated recursive feature elimination (RFE) procedure yields drastically different feature sets across software tools (see Figure 10). Overall all software tools require a small number of features for age prediction of control subjects (n*ϵ*[3, 20]) compared to ASD subjects (*nϵ*[41, 60]). RFE seems to be very sensitive to the QC procedures as these yield different feature sets with no apparent consistent trends for controls or ASD cohorts. The parcellation comparisons also show varied selection of features. Despite the larger number of regions for Destrieux and Glasser parcellations, the number of predictive features remain relatively small. The sensitivity to QC procedure appears to reflect in the parcellation analysis as evidenced by large spikes in feature counts for both control and ASD cohorts.

**Fig. 10.**
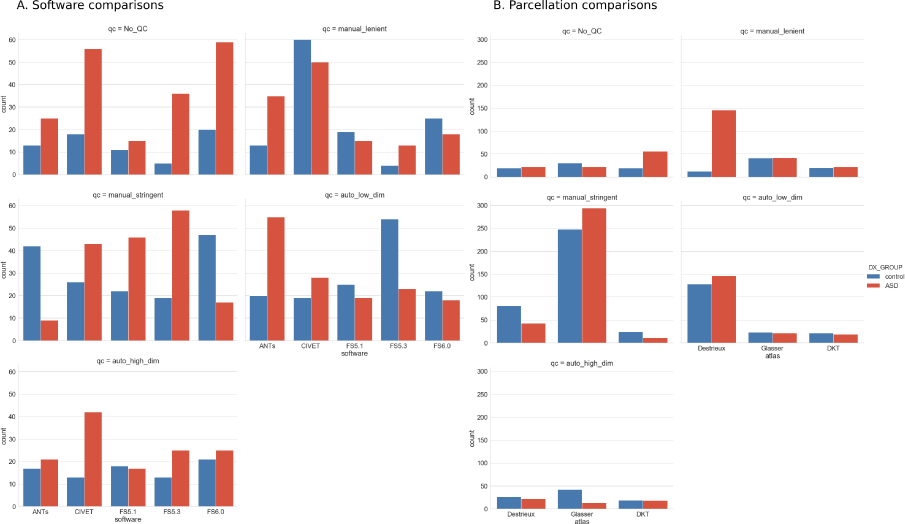
Predictive feature set count with various (A) software and (B) parcellations for different QC levels stratified by diagnosis. Optimal predictive features are selected using cross-validated recursive feature elimination procedure.

### Validation analysis

For the HCP dataset, the feature comparisons based on DKT parcellation yielded an average Pearson correlation of 0.66 between CIVET2.1 and FS6.0 (ABIDE: r=0.52). The regions exhibiting low correlations were also consistent with AIBIDE analysis, and comprised cingulate regions, orbitofrontal regions, entorhinal, pericalcarine, and insula.

## Discussion

In this work, we aimed to assess the reproducibility of phenotypic features and subsequent findings subjected to preprocessing pipeline variation along three axes: 1) image processing tool, 2) anatomical priors, 3) quality control. We emphasize that the goal here is not to deliberate specific biological and individual interpretation from the analyses, but rather to highlight the differences among the findings themselves, a key information for the large community of researchers using anatomical brain imaging in their studies.

In the TF-N analysis, we observe a weak ROI-wise correlation across software pairs (see Figure 3). Although software specific biases are expected in biological phenotypic estimates, the level of diminished correlation is striking. One can explain this performance for the comparisons involving ANTs as it is the only software that operates in the voxel (volume) space. However, a similarly poor performance is seen with CIVET and FreeSurfer, both of which operate in a vertex (surface) space for cortical thickness estimation. Since individual ROI-based measures are frequently used in the down-stream mass-univariate models, the lack of consensus across software tools is likely to yield different results. Moreover, the varying ROI covariance patterns across the software (see Figure. 4) suggest weak multivariate similarity, which again strongly increases the dependence of findings and biological interpretations on the software choice. For instance, the bilateral symmetry between cortical ROIs may only be inferred with CIVET due to its algorithmic specificities. Lastly, the lack of impact from QC suggests that these effects are systemic and not driven by outliers.

In the TF-I analysis, software tool specific t-SNE similarity is encouraging and expected. The t-SNE embeddings also highlight stronger differences between software tools compared to the differences in diagnostic groups (see Figure 5). This partly explains the high difficulty in training generalizable ML models across studies employing different preprocessing pipelines. The poor clustering consistency with commonly used Ward’s linkage criterion is alarming (see Table 4). Given that data-driven clustering is a typical practice to identify subgroups of patients or define meaningful biomarkers [44,45], clustering membership that is highly sensitive to the preprocessing pipeline may go undetected by the stability tests performed on the final set of processed features.

In the TD-N analysis, the software and parcellation comparisons show relatively consistent spatial associations for the age regression models (see Figure 7-8). There are some software-specific regional peculiarities (e.g. insular regions with CIVET), which also behave differently with various QC procedures as can be seen by more variable performance of ANTs and CIVET. These sensitivities should be noted as they could suggest methodological limitations or bias in the software. The overall cortex-wide association of thickness with age is expected as various studies have reported the same in healthy and ASD populations [38,40,46,47]. Direct comparison with other studies is challenging due to differences in the underlying statistical models, which produce varying topologies of wide-spread associations, and the direction of change in the cortical thickness. The results in this work suggest that the lack of strong ROI (univariate) correlation between a pair of software tools does not impact the task-driven mass-univariate analysis. However, we note that this is highly specific to the task at hand, as well as model selection procedures, which are beyond the scope of this work. We speculate that localized effects are likely to be more sensitive to the univariate pairwise relationships, and therefore a novel biological finding must be reported with high scrutiny to exclude pipeline specificities.

In the TD-I analysis, age prediction with random forest is stable subject to software and parcellation variations (see Figures 8-9). The RMSE performance of 3.8-4.7 years is comparable to the similar previous age prediction studies [16,37,38] that report RMSE in ranges of 6-12 years or mean absolute error of 1.7-1.95 years. The stability of performance could potentially be attributed to the relatively large sample sizes. It is encouraging to see that biological noise does not induce large variations into individual predictions. It is also important to note the impact of QC on the model performance and the null distributions for a given population (i.e. controls vs ASD). These alterations in the expected null performance need to be reported in order to fairly evaluate the improvements offered by a novel model on a given sample. Although random forest seems to be stable for individual predictions, the feature importance assessments by the same model are highly variable (see Figure 10). One explanation for this behaviour could be that in the presence of noisy biological features, ML models assign a relatively flat distribution of importance to the features. Variation in feature sets or sample sizes, as dictated by the selected preprocessing pipeline, would thus yield a drastically different feature ranking in a given iteration of the analysis. This needs to be taken into account if ML models are used to make biological inferences.

The validation analysis with HCP allowed us to replicate our feature correlations findings on an independent dataset. Similar to the ABIDE analysis, HCP data showed consistent low correlation between the ROI thickness values produced by FS6.0 and CIVET2.1. Moreover, there is a large overlap in the regions (i.e. cingulate regions, orbitofrontal regions, entorhinal, and insula) exhibiting the low correlations. This suggests that the low correlations are mainly driven by the algorithmic differences and not by the dataset. The pericalcarine was the exception to this common regional subset, which had a low correlation only in the HCP dataset, possibly due to dataset specific peculiarities. Nevertheless this highlights the need for larger meta-analyses to identify tool-specific and dataset-specific variability in findings.

### Limitations

Although in this work we aimed at assessing the impact of pipeline vibration along three different axes, we only considered a subset of permutations in the analysis. This was primarily due to practical reasons such as the lack of availability of common parcellation definitions for all software tools. Therefore we could not compare software tools with Destrieux and Glasser parcellations. We also limited the scope of this work to structural features, and did not consider functional or diffusion measures. With the increasing popularity of sophisticated, derived measures from highly flexible functional preprocessing pipelines with a multitude of design parameters, it is critical to understand and quantify the inherent variability and its impact on downstream findings. We defer this endeavor to future studies and refer to [6] for some progress in this direction.

## Conclusions

This work highlights the variability introduced by preprocessing pipelines, which is only a part of the larger issue of reproducibility in computational neuroimaging. We understand that the computational burden of comparative analyses such as described here can be infeasible in many studies. This necessitates undertaking of large meta analytic studies to understand software specific biases for various populations stratified by demographics and pathologies. At the single study level, we encourage the community to process data with different tools as much as possible and report variation of the results. We also propose to systematically report positive and negative results with different parcellations. This will improve confidence levels in the findings and help to better understand the spatial granularity associated with the effect of interest, while facilitating comparisons of common atlases across tools. Lastly, we also recommend assessing the sensitivity of findings against varying degrees of stringency for the QC criteria. Only with wide-spread adoption of rigorous scientific methodology and accessible informatics resources to replicate and compare processing pipelines can we address the compounding problem of reproducibility in the age of large-scale, data-driven computational neuroscience. The availability of containerized and well documented pipelines together with the necessary computing resources will mitigate the variability of results observed and direct the community towards understanding these differences, as well as further develop methodological validation and benchmarking.

## ACKNOWLEDGEMENTS

This work was partially funded by National Institutes of Health (NIH) NIH-NIBIB P41 EB019936 (ReproNim) NIH-NIMH R01 MH083320 (CANDIShare) and NIH RF1 MH120021 (NIDM), the National Institute Of Mental Health of the NIH under Award Number R01MH096906 (Neurosynth), as well as the Canada First Research Excellence Fund, awarded to McGill University for the Healthy Brains for Healthy Lives initiative and the Brain Canada Foundation with support from Health Canada. We thank Gleb Bezgin, John Lewis, and David Kennedy’s group for compiling manual QC lists used in this work. We are very grateful for a very thorough and insightful review of the manuscript by PJ Toussaint. We also thank Satrajit Ghosh for helping us extend this work as a stand-alone module for vibration analysis in future neuroimaging workflows.

## Supplementary information

Below are the validation results from task-free analyses on the HCP dataset. Figure 11 shows the regional correlations between CIVET2.1 and FS6.0 software. Figure 12 shows the t-SNE plot that highlights the software driven differences on individual clusters.

**Fig. 11.**
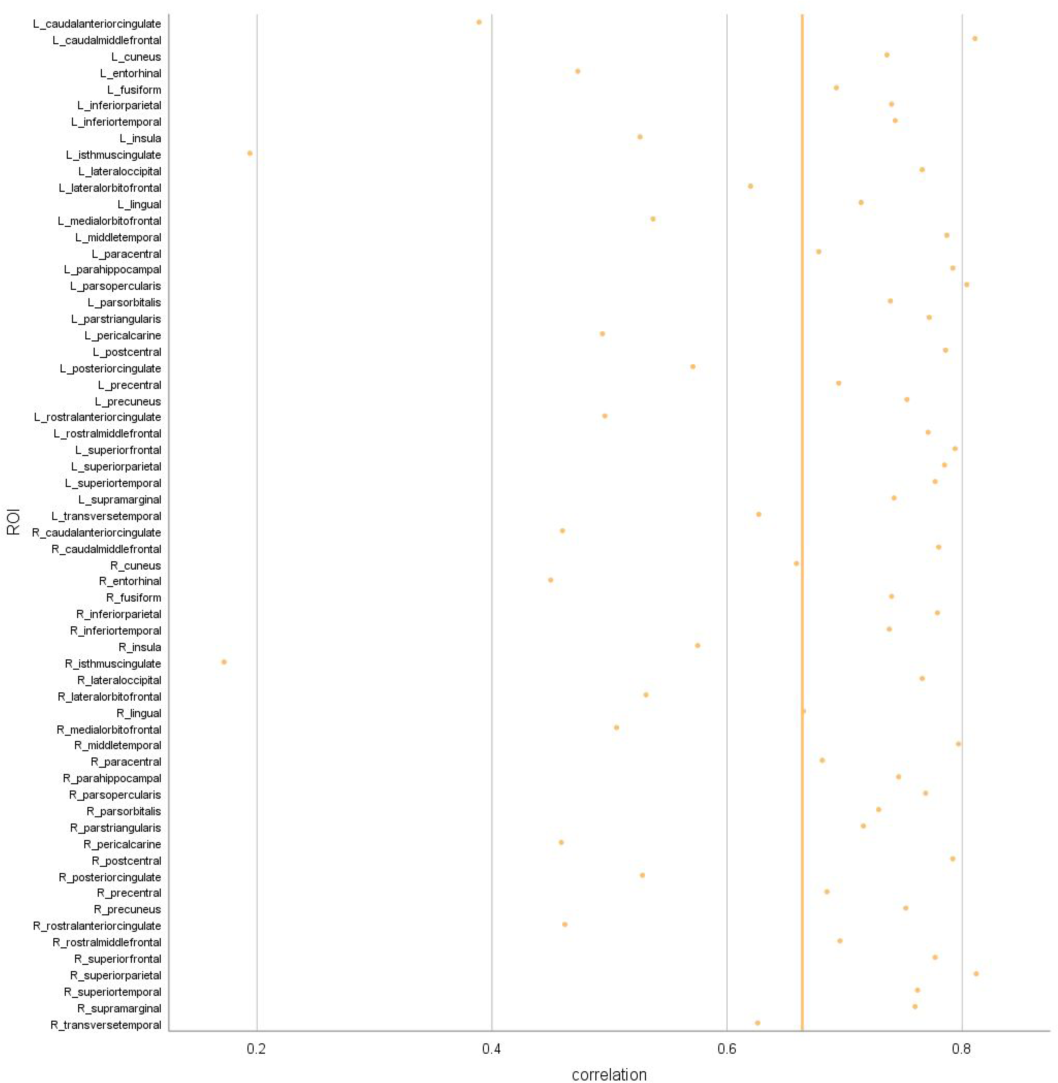
TF-N analysis for the HCP dataset: Left) Correlation between cortical thickness values for CIVET2.1 and FS6.0 measured independently over ROIs for control and ASD groups. The vertical lines represent the mean correlation across all ROIs, defined using DKT parcellation.

**Fig. 12.**
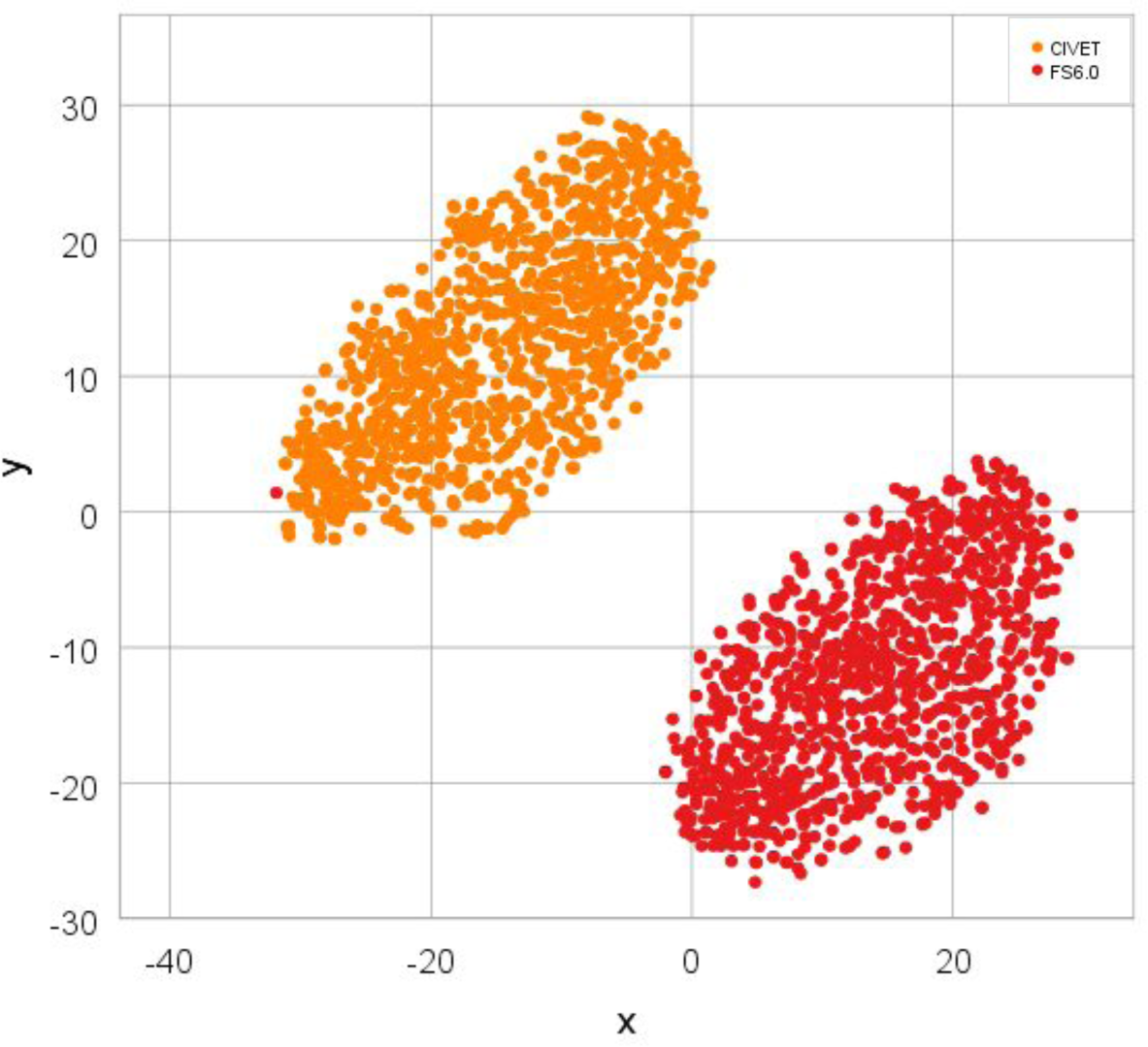
TF-N analysis for the HCP dataset: Left) Correlation between cortical thickness values for CIVET2.1 and FS6.0 measured independently over ROIs for control and ASD groups. The vertical lines represent the mean correlation across all ROIs, defined using DKT parcellation.

